# Using the UK Biobank as a global reference of worldwide populations: application to measuring ancestry diversity from GWAS summary statistics

**DOI:** 10.1101/2021.10.27.466078

**Authors:** Florian Privé

## Abstract

**Motivation:** Measuring genetic diversity is an important problem because increasing genetic diversity is key to making new genetic discoveries, while also being a major source of confounding to be aware of in genetics studies.

**Results:** Using the UK Biobank data, a prospective cohort study with deep genetic and phenotypic data collected on almost 500,000 individuals from across the United Kingdom, we carefully define 21 distinct ancestry groups from all four corners of the world. These ancestry groups can serve as a global reference of worldwide populations, with a handful of applications. Here we develop a method that uses allele frequencies and principal components derived from these ancestry groups to effectively measure ancestry proportions from allele frequencies of any genetic dataset.

**Availability:** This method is implemented as function snp_ancestry_summary as part of R package bigsnpr.

## Introduction

Several projects have focused on providing genetic data from diverse populations, such as the HapMap project, the 1000 genomes project (1KG), the Simons genome diversity project, and the human genome diversity project (International HapMap 3 Consortium *et al*. 2010; 1000 Genomes Project Consortium *et al*. 2015; Mallick *et al*. 2016; Bergström *et al*. 2020). However, these datasets do not contain many individuals per population and therefore are not large enough for some purposes, such as accurately estimating allele frequencies for diverse worldwide populations. The UK Biobank (UKBB) project is a prospective cohort study with deep genetic and phenotypic data collected on almost 500,000 individuals from across the United Kingdom. Despite being a cohort from the UK, this dataset is so large that it includes individuals that were born in all four corners of the world. Therefore, the UK Biobank can serve as a global reference of worldwide populations when used in its entirety, i.e. without discarding valuable multi-ancestry genetic data.

## Implementation

Here we carefully use information on self-reported ancestry, country of birth, and genetic similarity to define 21 distinct ancestry groups from the UK Biobank to be used as global reference populations, which is the first innovation of this paper. These include nine groups with genetic ancestries from Europe, four from Africa, three from South Asia, three from East Asia, one from the Middle East, and one from South America (which are later merged into 18 groups in table 1). The detailed procedure used to construct these reference ancestry groups is presented in the Supplementary Materials. As a direct application of these groups, we propose a new method to estimate global ancestry proportions from a cohort based on its allele frequencies only (i.e. summary statistics). Arriaga-MacKenzie *et al*. (2021) previously proposed method Summix, which finds the convex combination of ancestry proportions *α*_*k*_ (positive and sum to 1) which minimizes the following problem: 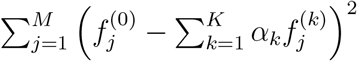, where *M* is the number of variants, *K* the number of reference populations, 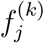 is the frequency of variant *j* in population *k*, and 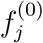 is the frequency of variant *j* in the cohort of interest. Arriaga-MacKenzie *et al*. (2021) used the five continental 1KG populations as reference.

**Table 1:**
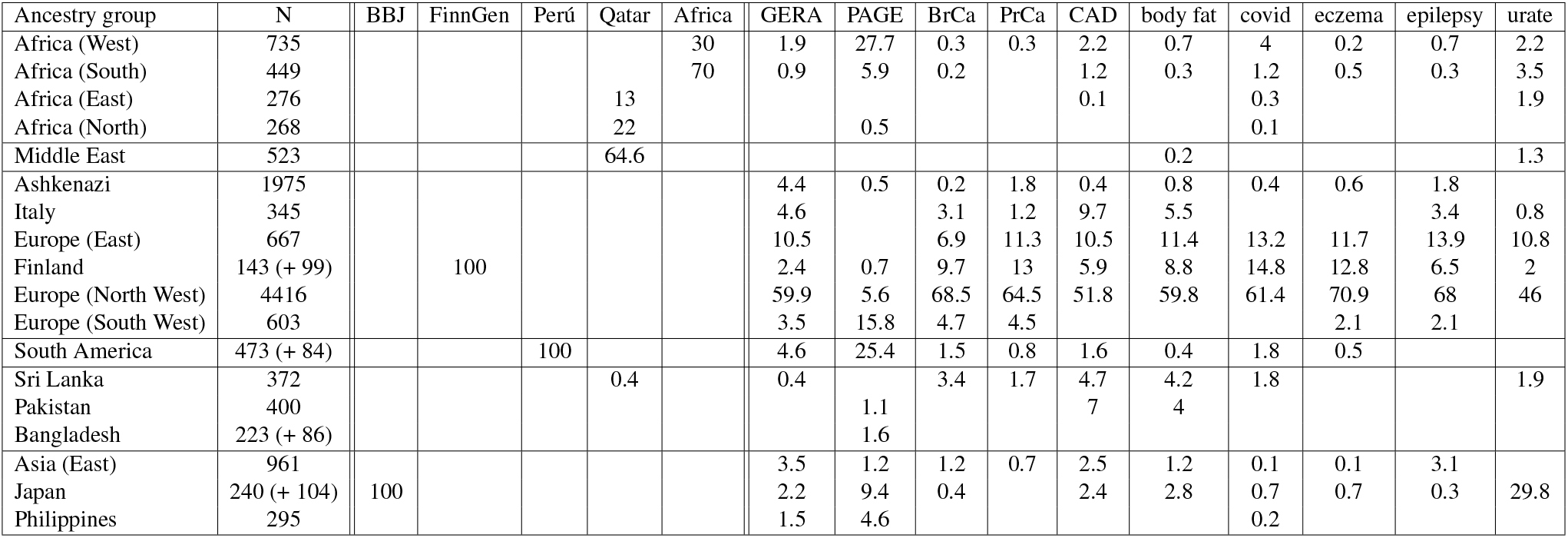
Reference populations with their size (N), and corresponding ancestry proportions (in %) inferred from the proposed `snp_ancestry_summary` method, for several GWAS summary statistics. Note that, because they are very close ancestry groups, we merge a posteriori the ancestry coefficients *α*_*k*_ from “Ireland”, “United Kingdom” and “Scandinavia” into a single “Europe (North West)” group, and similarly for “Europe (North East)” and “Europe (South East)” into a single “Europe (East)” group. Citations for the allele frequencies used: the Biobank Japan (BBJ, Sakaue *et al*. (2021)), FinnGen (https://r5.finngen.fi/), GWAS in Peruvians (Asgari *et al*. 2020), GWAS in Qataris (Thareja *et al*. 2021), GWAS in Sub-Saharan Africans (Africa, Chen *et al*. (2019)), GERA (Hoffmann *et al*. 2018), PAGE (Wojcik *et al*. 2019), breast cancer (BrCa, Michailidou *et al*. (2017)), prostate cancer (PrCa, Schu-macher *et al*. (2018)), coronary artery disease (CAD, Nikpay *et al*. (2015)), body fat percentage (Lu *et al*. 2016), COVID-19 (The COVID-19 Host Genetics Initiative *et al*. 2021), eczema (Paternoster *et al*. 2015), epilepsy (The International League Against Epilepsy Consortium on Complex Epilepsies 2018), and serum urate (Tin *et al*. 2019). Several of these GWAS summary statistics have been downloaded through the NHGRI-EBI GWAS Catalog (MacArthur *et al*. 2017).

| Ancestry group | N | BBJ | FinnGen | Perú | Qatar | Africa | GERA | PAGE | BrCa | PrCa | CAD | body fat | covid | eczema | epilepsy | urate |
| --- | --- | --- | --- | --- | --- | --- | --- | --- | --- | --- | --- | --- | --- | --- | --- | --- |
| Africa (West) | 735 |  |  |  |  | 30 | 1.9 | 27.7 | 0.3 | 0.3 | 2.2 | 0.7 | 4 | 0.2 | 0.7 | 2.2 |
| Africa (South) | 449 |  |  |  |  | 70 | 0.9 | 5.9 | 0.2 |  | 1.2 | 0.3 | 1.2 | 0.5 | 0.3 | 3.5 |
| Africa (East) | 276 |  |  |  | 13 |  |  |  |  |  | 0.1 |  | 0.3 |  |  | 1.9 |
| Africa (North) | 268 |  |  |  | 22 |  |  | 0.5 |  |  |  |  | 0.1 |  |  |  |
| Middle East | 523 |  |  |  | 64.6 |  |  |  |  |  |  | 0.2 |  |  |  | 1.3 |
| Ashkenazi | 1975 |  |  |  |  |  | 4.4 | 0.5 | 0.2 | 1.8 | 0.4 | 0.8 | 0.4 | 0.6 | 1.8 |  |
| Italy | 345 |  |  |  |  |  | 4.6 |  | 3.1 | 1.2 | 9.7 | 5.5 |  |  | 3.4 | 0.8 |
| Europe (East) | 667 |  |  |  |  |  | 10.5 |  | 6.9 | 11.3 | 10.5 | 11.4 | 13.2 | 11.7 | 13.9 | 10.8 |
| Finland | 143 (+ 99) |  | 100 |  |  |  | 2.4 | 0.7 | 9.7 | 13 | 5.9 | 8.8 | 14.8 | 12.8 | 6.5 | 2 |
| Europe (North West) | 4416 |  |  |  |  |  | 59.9 | 5.6 | 68.5 | 64.5 | 51.8 | 59.8 | 61.4 | 70.9 | 68 | 46 |
| Europe (South West) | 603 |  |  |  |  |  | 3.5 | 15.8 | 4.7 | 4.5 |  |  |  | 2.1 | 2.1 |  |
| South America | 473 (+ 84) |  |  | 100 |  |  | 4.6 | 25.4 | 1.5 | 0.8 | 1.6 | 0.4 | 1.8 | 0.5 |  |  |
| Sri Lanka | 372 |  |  |  | 0.4 |  | 0.4 |  | 3.4 | 1.7 | 4.7 | 4.2 | 1.8 |  |  | 1.9 |
| Pakistan | 400 |  |  |  |  |  |  | 1.1 |  |  | 7 | 4 |  |  |  |  |
| Bangladesh | 223 (+ 86) |  |  |  |  |  |  | 1.6 |  |  |  |  |  |  |  |  |
| Asia (East) | 961 |  |  |  |  |  | 3.5 | 1.2 | 1.2 | 0.7 | 2.5 | 1.2 | 0.1 | 0.1 | 3.1 |  |
| Japan | 240 (+ 104) | 100 |  |  |  |  | 2.2 | 9.4 | 0.4 |  | 2.4 | 2.8 | 0.7 | 0.7 | 0.3 | 29.8 |
| Philippines | 295 |  |  |  |  |  | 1.5 | 4.6 |  |  |  |  | 0.2 |  |  |  |

Here we provide reference allele frequencies for 5,816,590 genetic variants across 21 diverse ancestry groups (which are later merged into 18 groups in table 1). Moreover, we rely on the projection of our reference allele frequencies onto the PCA (principal component analysis) space computed from the corresponding UKBB (and 1KG) individuals, and also make these PC loadings available for download. Instead, we then minimize 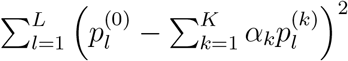, with the same convex constraints on ancestry proportions *α*_*k*_, and where *L* is the number of PCs, 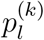 is the projection of allele frequencies from population *k* onto PC *l*, and 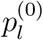 is the (corrected) projection of allele frequencies from the cohort of interest onto PC *l*. Note that we need to correct for the shrinkage when projecting a new dataset (here the allele frequencies from the GWAS summary statistics) onto the PC space (Privé *et al*. 2020). Finding the ancestry proportions in the PCA space (rather than using the allele frequencies directly) provides more power to distinguish between close populations, which is the second innovation of this paper. This enables us to use more reference populations in order to get a more fine-grained measure of genetic diversity.

The steps required by the proposed method are then 1/ read all summary statistics datasets into R, i.e. the reference allele frequencies and corresponding PC loadings we provide for download as well as the GWAS summary statistics containing the allele frequencies of interest; 2/ match variants and alleles between summary statistics and the reference allele frequencies we provide; 3/ project allele frequencies onto the PCA space (matrix multiplication); 4/ solve the final (small) quadratic programming problem, by relying on R package quadprog (Turlach *et al*. 2019). Steps 3 and 4 are now implemented in function `snp_ancestry_summary` in our R package bigsnpr (Privé *et al*. 2018). Step 2 can be performed using existing function `snp_match`. A tutorial is provided at https://privefl.github.io/bigsnpr/articles/ancestry.html. All these steps are very fast and overall require a few minutes only for GWAS summary statistics with millions of variants.

## Results

We download several GWAS summary statistics for which allele frequencies are reported, and apply this new method to them. We first apply function `snp_ancestry_summary` to more homogeneous samples as an empirical validation; when applying it to the Biobank Japan (Japanese cohort), FinnGen (Finnish), a Peruvian cohort, a Qatari cohort and Sub-Saharan African cohort, the ancestry proportions obtained match expectations (Table 1). When comparing our estimates with reported ancestries for more diverse cohorts, for example PAGE is composed of 44.6% Hispanic-Latinos, 34.7% African-Americans, 9.4% Asians, 7.9% Native Hawaiians and 3.4% of some other ancestries (self-reported), whereas our estimates are of 25.4% South American, 22.6% European (including 15.8% from South-West Europe), 34.1% African, 2.7% South Asian, 10.6% East Asian, and 4.6% Filipino. GWAS summary statistics from either European ancestries or more diverse ancestries all have a substantial proportion estimated from European ancestry groups, while ancestries from other continents are still largely underrepresented (Table 1).

We then perform three secondary analyses. First, we compare the results obtained previously in Table 1 with the results we would get without using the PCA projection of allele frequencies (i.e. equivalent to the Summix method). The resulting ancestry proportions are presented in Table S1 and are clearly less precise for Biobank Japan (BBJ) and FinnGen. Second, we compare previous results with the ones obtained using a smaller number of variants, by randomly sampling 100,000 variants to run the proposed method. The resulting ancestry proportions are presented in Table S2 and are highly consistent with the ones from Table 1, showing that 100,000 overlapping variants are enough to run the proposed method.

Third, we also infer ancestry proportions for all 345 individuals of the Simons genome diversity project (Mallick *et al*. 2016) using the reference allele frequencies we provide and two methods. We use either our proposed method with the genotypes of an individual divided by 2 in place of allele frequencies, or by using the projection analysis of ADMIXTURE (`-P`, Shringarpure *et al*. (2016)). Results are very consistent between the two methods, and are overall as expected, further validating the proposed ancestry groups and the proposed method to infer ancestry proportions, which seems very precise even at the individual level.

## Discussion

Here we have identified an unprecedentedly large and diverse set of ancestry groups within a single cohort, the UK Biobank. Using allele frequencies and principal components (PCs) derived from these ancestry groups, we show how to effectively measure diversity from GWAS summary statistics reporting allele frequencies. Measuring genetic diversity is an important problem because increasing genetic diversity is key to making new genetic discoveries, while also being a major source of confounding to be aware of in genetics studies. Our work has limitations though. First, it is unknown whether we can effectively capture any existing ancestry as a combination of the 21 reference populations we defined. For example, it seems that Native Hawaiians in the PAGE study are partly captured by the “Philippines” ancestry group we define. Second, with the 21 ancestry groups we define, we probably capture a large proportion of the genetic diversity in Europe, but more fine-grained diversity in other continents may still be lacking. Third, when using the allele frequencies reported in the GWAS summary statistics, it is not clear whether they were computed from all individuals (i.e. before performing any quality control and filtering), and, for meta-analyses of binary traits, whether they were computed as a weighted average of total or effective sample sizes. Despite these limitations, we envision that the ancestry groups we define here will have many useful applications. The presented method that uses these groups could e.g. be used to automatically report ancestry proportions in the GWAS Catalog (MacArthur *et al*. 2017). These ancestry groups could also be used for assigning ancestry in other cohorts using the PC projection from this study (Privé *et al*. 2022), assessing portability of polygenic scores (Privé *et al*. 2022), or deriving linkage disequilibrium (LD) references matching GWAS summary statistics from diverse ancestries.

## Software and code availability

The newest version of R package bigsnpr can be installed from GitHub (see https://github.com/privefl/bigsnpr) and a recent enough version can be installed from CRAN. A tutorial on ancestry proportions and ancestry grouping is available at https://privefl.github.io/bigsnpr/articles/ancestry.html. The set of reference allele frequencies for 5,816,590 genetic variants across 21 diverse ancestry groups defined here can be downloaded at https://figshare.com/ndownloader/files/31620968 and PC loadings for all variants across 16 PCs at https://figshare.com/ndownloader/files/31620953. All code used for this paper is available at https://github.com/privefl/freq-ancestry/tree/main/code. We have extensively used R packages bigstatsr and bigsnpr (Privé *et al*. 2018) for analyzing large genetic data, packages from the future framework (Bengtsson 2021) for easy scheduling and parallelization of analyses on the HPC cluster, and packages from the tidyverse suite (Wickham *et al*. 2019) for shaping and visualizing results.

## Acknowledgements

F.P. is supported by the Danish National Research Foundation (Niels Bohr Professorship to John McGrath) and by a Lundbeck Foundation Fellowship (R335-2019-2339 to Bjarni J. Vilhjálmsson). The author thanks Shai Carmi for helpful discussions. The author thanks GenomeDK and Aarhus University for providing computational resources and support that contributed to these research results. This research has been conducted using the UK Biobank Resource under Application Number 58024.

