## Supplementary Materials for "Using the UK Biobank as a global reference of worldwide populations: application to measuring ancestry diversity from GWAS summary statistics"

### S1 Designing a reference of allele frequencies from the UK Biobank

#### S1.1 Identifying 26 ancestry groups from the UK Biobank

In this section, we use the principal components (PCs, Data-Field 22009), the self-reported ancestry (Data-Field 21000), and the country of birth (Data-Field 20115) from the UK Biobank (Bycroft *et al.* 2018) to first identify 26 homogeneous ancestry groups. We use the first 16 PCs only, which we have shown to be the PCs capturing population structure in the UK Biobank (Privé *et al.* 2020). We compute Euclidean distances (L2-norms) from these PCs (i.e. in the PCA space), which we have shown to be an appropriate measure of the genetic distance between populations since these (squared) distances are largely proportional to the $F_{ST}$ between populations (Privé *et al.* 2022).

For the individuals with a missing country of birth, we assign to them "United Kingdom" when they self-identify as "British" and "Ireland" when they self-identify as "Irish". Since the country of birth may poorly corresponds to the genetic ancestry that we are interested in, we perform some quality control on this information. Indeed, there are many individuals born in Africa with an Asian self-reported ancestry; we set their country of birth to missing. We also set this information to missing for 1/ individuals born outside of Europe that are too close to the UK (log-distance to the UK center lower than 4); 2/ individuals born outside the UK within a log-distance of 2.5 to the UK; 3/ individuals born in Europe not within a log-distance of 5 to the UK. Finally, for individuals born in South Africa, we keep only the ones with a log-distance to the UK larger than 6 (individuals of African ancestry). Note, for reference, that Ireland is at a log-distance of 3.3 from the UK, the Middle East at 4.7 and West Africa at 6. After applying these filters, there are still many individuals born outside of the UK and Ireland, including e.g. 3352 individuals from India, 1100 from Germany, 1036 from Nigeria, and 76 countries with at least 50 individuals.

Then, to construct a homogeneous ancestry group to be used as a reference population, we choose a country, compute the robust PC center of individuals born in this country (using the geometric median, Privé *et al.* (2022)), and choose to include in this group all individuals within a chosen log-distance to the center. We investigate multiple choices for the countries and distance thresholds by carefully and manually inspecting the self-reported ancestry and country of birth of the individuals kept or excluded from these groups. For groups named after a country, we further restrict to individuals born in this country. For the United Kingdom and Ireland groups, we pick 2000 individuals at random for each, in order not to have very uneven (larger) sample sizes compared to the other groups. We manually investigate different country centers and distance thresholds to cover as much of the worldwide populations as possible, without having overlapping groups. We also include two other groups: one "Ashkenazi" group based on a PC center we defined in Privé *et al.* (2022), and a "South America" group based on individuals with PC6 $> 50$ and PC8 $< -30$ (clearly separated from the rest of individuals from the UK Biobank). Finally, we investigate individuals that are far away from any of the groups defined previously, and add a last group based on a center from remaining individuals born in India.

For each of the 26 ancestry groups we define, we list the individuals' countries of birth (with at least 5 individuals, where "NA" means unknown):

- **Japan:** Japan: 240
- **Asia (East):** Hong Kong: 390 — Malaysia: 179 — NA: 122 — China: 114 — Singapore: 58 — Vietnam: 33 — Indonesia: 14 — Taiwan: 10 — Thailand: 8 — Brunei: 6 — Macau (Macao): 6
- **Philippines:** Philippines: 295
- **Africa (South):** Zimbabwe: 251 — Uganda: 57 — South Africa: 47 — Zambia: 41 — Malawi: 10 — NA: 10 — Tanzania: 6 — Congo: 5
- **Africa (North):** Algeria: 69 — Morocco: 62 — Egypt: 54 — Libya: 36 — NA: 18 — Tunisia: 13
- **Africa (East 1):** Somalia: 81 — Ethiopia: 58 — Sudan: 54 — Eritrea: 46 — NA: 27
- **Africa (East 2):** Kenya: 104 — Uganda: 60 — Burundi: 18 — NA: 16 — Sudan: 14 — Tanzania: 14 — Rwanda: 13 — Nigeria: 8 — Somalia: 7 — Angola: 5 — Zimbabwe: 5
- **Caribbean:** Caribbean: 327 — NA: 310 — Barbados: 73 — The Guianas: 23 — Antigua and Barbuda: 7
- **Africa (West):** Nigeria: 217 — Ghana: 175 — NA: 122 — Sierra Leone: 98 — Caribbean: 80 — Ivory Coast: 7 — Liberia: 7 — The Guianas: 7 — Togo: 6
- **Africa (Central):** Congo: 117 — Cameroon: 42 — Nigeria: 33 — NA: 33 — Angola: 21 — Caribbean: 8 — The Guianas: 7
- **Middle East:** Iraq: 240 — Iran: 172 — Turkey: 55 — NA: 31 — Syria: 11
- **United Kingdom:** United Kingdom: 2000
- **Ireland:** Ireland: 2000

- **Finland:** Finland: 143
- **Scandinavia:** Denmark: 142 — Norway: 82 — Sweden: 75 — NA: 68 — Germany: 25 — Netherlands: 17 — Iceland: 5
- **Europe (South West):** Spain: 279 — Portugal: 198 — NA: 91 — France: 25 — Gibraltar: 7
- **Italy:** Italy: 345
- **Europe (South East):** NA: 115 — Romania: 47 — Serbia/Montenegro: 41 — Bosnia and Herzegovina: 34 — Bulgaria: 33 — Croatia: 28 — Hungary: 17 — Italy: 11 — Macedonia: 8
- **Europe (Central):** NA: 196 — Germany: 111 — Czech Republic: 79 — Poland: 50 — Austria: 33 — Hungary: 27 — Slovakia: 25 — Croatia: 10 — Slovenia: 8 — France: 7
- **Europe (North East):** Russia: 88 — NA: 87 — Poland: 72 — Ukraine: 34 — Latvia: 10 — Germany: 7 — Kazakhstan: 6 — Lithuania: 5
- **Pakistan:** Pakistan: 400
- **Sri Lanka:** Sri Lanka: 372
- **Bangladesh:** Bangladesh: 223
- **South America:** Colombia: 173 — Chile: 57 — Mexico: 51 — Peru: 50 — Ecuador: 33 — Bolivia: 21 — NA: 20 — Venezuela: 15 — Brazil: 10 — United Kingdom: 8 — USA: 8 — Argentina: 5
- **Ashkenazi:** United Kingdom: 1182 — NA: 572 — USA: 115 — Israel: 27 — Hungary: 12 — Ireland: 8 — France: 7 — Canada: 6 — Russia: 5
- **India:** NA: 258 — India: 108 — Sri Lanka: 16 — Pakistan: 11

### S1.2 Computing allele frequencies

We download the 1000 Genomes (1KG) Project data (1000 Genomes Project Consortium *et al.* 2015). Using PLINK (Chang *et al.* 2015), we remove 9 outlier individuals identified in Martin *et al.* (2017), filter for founders, non-sex chromosomes, non-multiallelic variants with minor allele counts of at least 10. From the UK Biobank, we select imputed variants with a minor allele frequency larger than 0.001 and INFO score larger than 0.3, and match these variants with the ones from the filtered 1KG data described previously. We identify 12,373,666 genetic variants in common, for which we compute allele frequencies for each of the 26 1KG populations. We also compute allele frequencies from the BGEN imputed data for each of the 26 UK Biobank groups we identified previously.

### S1.3 Designing the final set of variants

We select variants with an INFO score of at least 0.4 in all 26 UK Biobank ancestry groups. Based on the remaining 5,840,630 variants, we compute the overall fixation indices $F_{ST}$ between the UK Biobank groups and the 1KG populations (See the description of the 1KG populations at `https://www.coriell.org/1/NHGRI/Collections/1000-Genomes-Collections/1000-Genomes-Project)`. We report all $F_{ST}$ at `https://github.com/privefl/freq-ancestry/blob/main/all_fst.csv`, and show only the lowest for each of the 26 ancestry groups we define here:

- **Japan:** JPT: 0.00033 — CHB: 0.007 — CHS: 0.0089 — Asia (East): 0.01 — KHV: 0.014 — CDX: 0.017 — Philippines: 0.021
- **Asia (East):** CHS: 0.0009 — KHV: 0.0021 — CHB: 0.0029 — CDX: 0.0034 — Philippines: 0.01 — Japan: 0.01 — JPT: 0.011
- **Philippines:** Asia (East): 0.01 — KHV: 0.01 — CDX: 0.011 — CHS: 0.012 — CHB: 0.015 — Japan: 0.021 — JPT: 0.021
- **Africa (South):** Africa (Central): 0.0017 — LWK: 0.0035 — Caribbean: 0.0043 — Africa (West): 0.0044 — Africa (East 2): 0.005 — YRI: 0.005 — ESN: 0.0055
- **Africa (North):** PUR: 0.01 — Middle East: 0.01 — Italy: 0.01 — Europe (South West): 0.011 — TSI: 0.012 — IBS: 0.012 — Ashkenazi: 0.012
- **Africa (East 1):** Africa (North): 0.018 — ASW: 0.027 — Africa (East 2): 0.028 — PUR: 0.029 — Middle East: 0.036 — ACB: 0.037 — CLM: 0.039
- **Africa (East 2):** LWK: 0.0033 — Caribbean: 0.0045 — ASW: 0.0048 — ACB: 0.0048 — Africa (South): 0.005 — Africa (Central): 0.0056 — Africa (West): 0.0074
- **Caribbean:** ACB: 0.00048 — Africa (West): 0.0011 — YRI: 0.0022 — Africa (Central): 0.0023 — ESN: 0.0028 — ASW: 0.003 — MSL: 0.0042
- **Africa (West):** YRI: 0.00061 — Caribbean: 0.0011 — ESN: 0.0015 — Africa (Central): 0.0019 — ACB: 0.0023 — MSL: 0.0028 — Africa (South): 0.0044
- **Africa (Central):** Africa (South): 0.0017 — Africa (West): 0.0019 — Caribbean: 0.0023 — YRI: 0.0024 — ESN: 0.0028 — ACB: 0.0036 — LWK: 0.0045
- **Middle East:** Italy: 0.0043 — TSI: 0.0059 — Europe (South East): 0.007 — Ashkenazi: 0.007 — Europe (South West): 0.008 — India: 0.0088 — IBS: 0.0089
- **United Kingdom:** Scandinavia: 0.00043 — Ireland: 0.00069 — CEU: 0.001 — GBR: 0.0011 — Europe (Central): 0.0012 — Europe (South East): 0.0021 — Europe (South West): 0.0023

- **Ireland:** United Kingdom: 0.00069 — Scandinavia: 0.0015 — GBR: 0.0017 — CEU: 0.0018 — Europe (Central): 0.0023 — Europe (South West): 0.0033 — Europe (South East): 0.0033
- **Finland:** FIN: 0.00041 — Europe (North East): 0.0052 — Scandinavia: 0.0054 — Europe (Central): 0.0056 — United Kingdom: 0.0066 — CEU: 0.0067 — GBR: 0.0072
- **Scandinavia:** United Kingdom: 0.00043 — Europe (Central): 0.0012 — CEU: 0.0013 — Ireland: 0.0015 — GBR: 0.0015 — Europe (South East): 0.0026 — Europe (North East): 0.0027
- **Europe (South West):** IBS: 0.00079 — Italy: 0.0014 — Europe (South East): 0.002 — TSI: 0.0021 — United Kingdom: 0.0023 — Europe (Central): 0.003 — CEU: 0.003
- **Italy:** TSI: 0.0011 — Europe (South West): 0.0014 — Europe (South East): 0.0017 — IBS: 0.0023 — Europe (Central): 0.004 — United Kingdom: 0.0041 — Middle East: 0.0043
- **Europe (South East):** Europe (Central): 0.00075 — Italy: 0.0017 — Europe (North East): 0.0017 — Europe (South West): 0.002 — United Kingdom: 0.0021 — Scandinavia: 0.0026 — TSI: 0.0027
- **Europe (Central):** Europe (North East): 0.00053 — Europe (South East): 0.00075 — Scandinavia: 0.0012 — United Kingdom: 0.0012 — CEU: 0.0019 — GBR: 0.0023 — Ireland: 0.0023
- **Europe (North East):** Europe (Central): 0.00053 — Europe (South East): 0.0017 — Scandinavia: 0.0027 — United Kingdom: 0.003 — CEU: 0.0036 — Ireland: 0.0039 — GBR: 0.0039
- **Pakistan:** PJL: 0.0026 — India: 0.0041 — GIH: 0.0053 — Bangladesh: 0.0059 — Sri Lanka: 0.0065 — ITU: 0.0068 — STU: 0.0074
- **Sri Lanka:** STU: 0.00034 — ITU: 0.0015 — Bangladesh: 0.0023 — BEB: 0.0025 — PJL: 0.0037 — GIH: 0.0043 — Pakistan: 0.0065
- **Bangladesh:** BEB: 0.00068 — Sri Lanka: 0.0023 — STU: 0.0028 — ITU: 0.003 — PJL: 0.0037 — GIH: 0.0044 — Pakistan: 0.0059
- **South America:** MXL: 0.0023 — CLM: 0.0053 — PUR: 0.014 — PEL: 0.019 — India: 0.022 — Pakistan: 0.027 — Europe (South East): 0.029
- **Ashkenazi:** Italy: 0.0051 — TSI: 0.0064 — Europe (South West): 0.0069 — Middle East: 0.007 — Europe (South East): 0.0072 — IBS: 0.008 — Europe (Central): 0.0097
- **India:** Pakistan: 0.0041 — PJL: 0.007 — Middle East: 0.0088 — Europe (South East): 0.0091 — Bangladesh: 0.0095 — Europe (Central): 0.0096 — United Kingdom: 0.0097

For each of the 1KG populations, we identify the closest ancestry group among those we defined, compute the $F_{ST}$ for all variants and scale them by dividing by the overall $F_{ST}$ with this group. For each variant, we then sum these scaled $F_{ST}$ (one for each 1KG population), and identify and remove 24,040 variants with a large score. This enable us to identify variants with possible errors in the UK Biobank. We obtain a set of 5,816,590 variants left.

### S1.4 Designing the final set of reference populations and preparing PCA projection

The 26 ancestry groups defined previously comprise a total of 15,645 individuals from the UK Biobank. We combine them with 2490 individuals from the 26 populations of the 1KG data, and use a pruned set of 252,893 high-quality genotyped variants to run an admixture analysis (K = 15) using function `snmf` from R package LEA (Frichot *et al.* 2014; Gain and François 2020). Ancestry proportions for the 52 groups are represented in figure S1. We can see that we have defined very homogeneous groups. Then, based on these results and previous $F_{ST}$, we merge four of the groups we defined with some 1KG populations. Indeed, for the "Finland", "Bangladesh" and "Japan" ancestry groups, we compute a weighted average of allele frequencies with the ones from the 1KG data (FIN, BEB and JPT) to increase the small sample size of these groups. We do the same for "South America" and "PEL" in order to capture slightly more of what is probably an Amerindian ancestral component (in blue).

We then compute the PCA from the 15,645 UKBB individuals and the 1KG individuals from the four added populations, across the 5,816,590 variants. We do not center the genotype matrix to capture one more component, and for an easier projection later. However, we do scale the genotypes by dividing them by the square root of LD scores to prevent from capturing LD structure in the PCA (Zou *et al.* 2010). We verify that PC loadings are homogeneously distributed along the genome (no peak) in figure S2 and that PC scores are capturing population structure in figure S3 (Privé *et al.* 2020). Based on the PC scores and admixture results (Figures S1 and S3), we remove some groups that are admixed or too central, which therefore could be retrieved from combining two or three other groups: "Africa (East 2)", "Caribbean", "Africa (Central)", "Europe (Central)", and "India". This results in a set of 5,816,590 variants for 21 ancestry groups, which we provide for download and use to estimate ancestry proportions in GWAS summary statistics. We also provide PC loadings scaled so that they can be directly used with allele frequencies. Note that, due to the shrinkage when projecting PCs, a correction needs to be applied when projecting new allele frequencies (Privé *et al.* 2020). We use six 1KG populations (YRI, IBS, STU, CHS, GBR, and TSI) and the corresponding UKBB ancestry groups we defined ("Africa (West)", "Europe (South West)", "Sri Lanka", "Asia (East)", "United Kingdom", and "Italy") to estimate the correction coefficients. From PC1 to PC16, these are 1.000, 1.000, 1.000, 1.008, 1.021, 1.034, 1.052, 1.074, 1.099, 1.123, 1.150, 1.195, 1.256, 1.321, 1.382, and 1.443.

Finally, the convex combination of ancestry proportions $\alpha_k$ (positive and sum to 1) is estimated by minimizing the following problem: $\sum_{l=1}^{L} \left( p_l^{(0)} - \sum_{k=1}^{K} \alpha_k p_l^{(k)} \right)^2$, where $L$ is the number of PCs (16

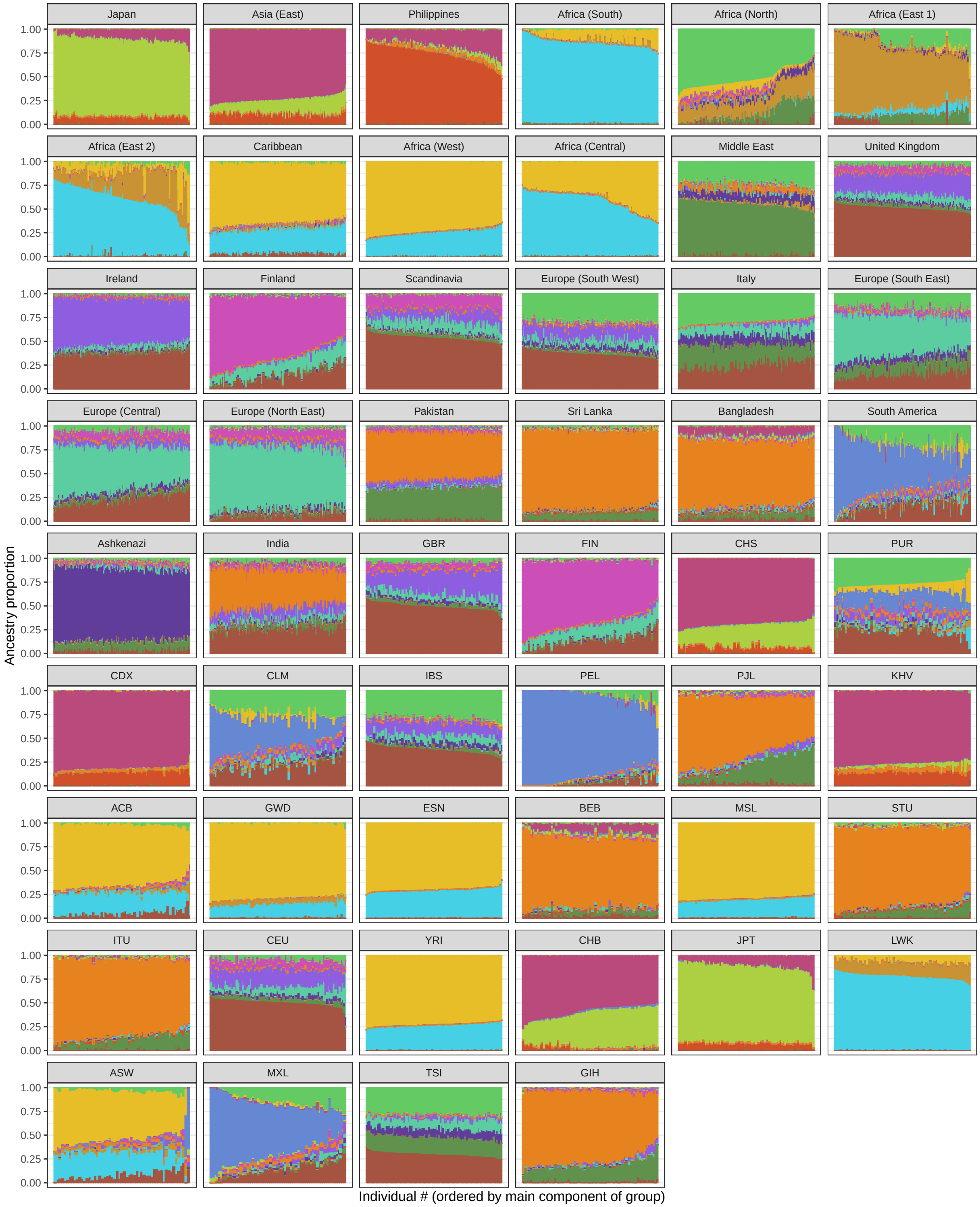


Figure S1: Admixture coefficients (for K = 15 ancestral populations) for the individuals from the 26 ancestry groups we defined previously from the UK Biobank, as well as the 26 1KG populations.

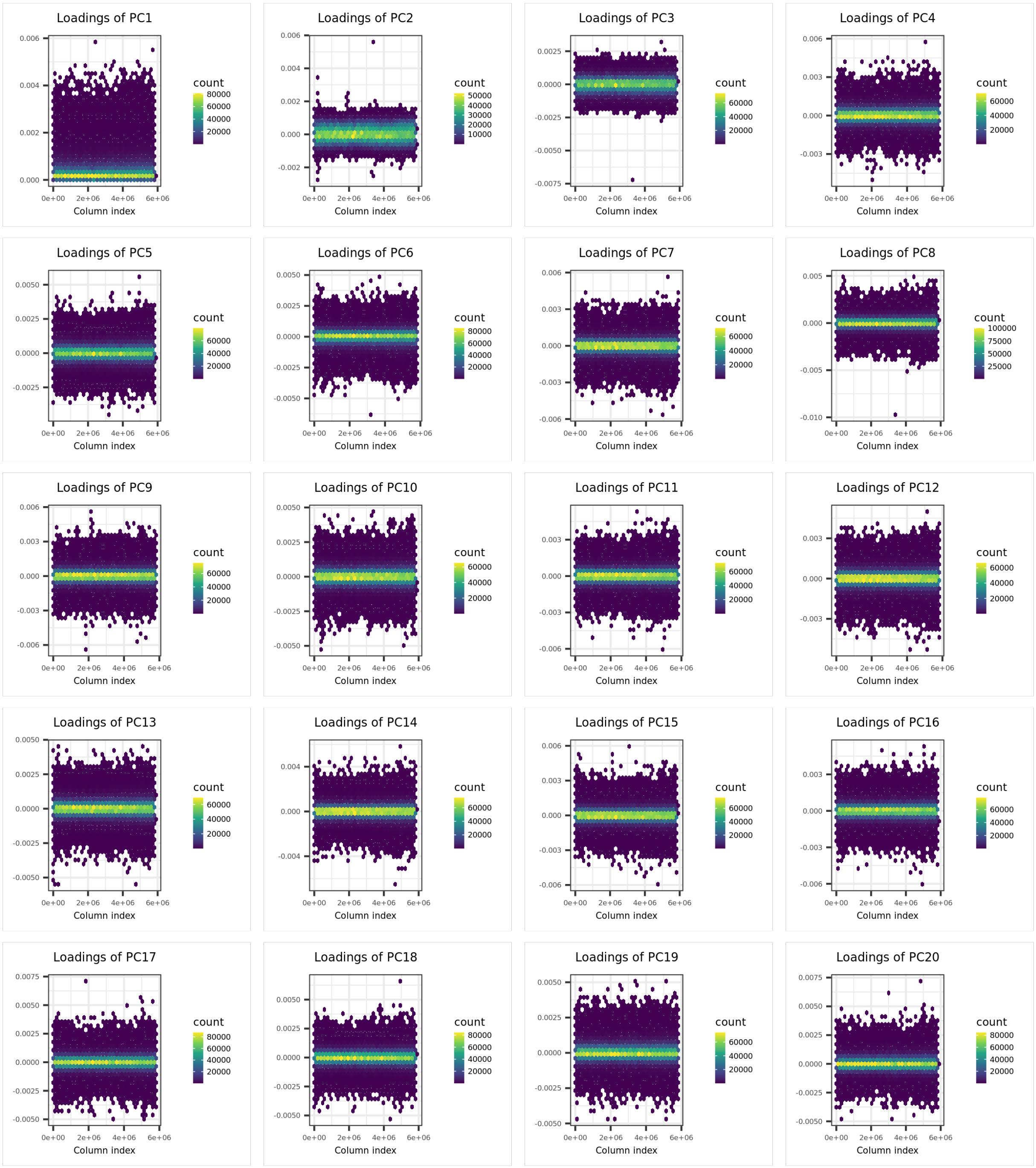


Figure S2: PCA loadings over 5.8M variants for the PCA recomputed from the subset of UKBB and 1KG individuals used in the reference ancestry groups.

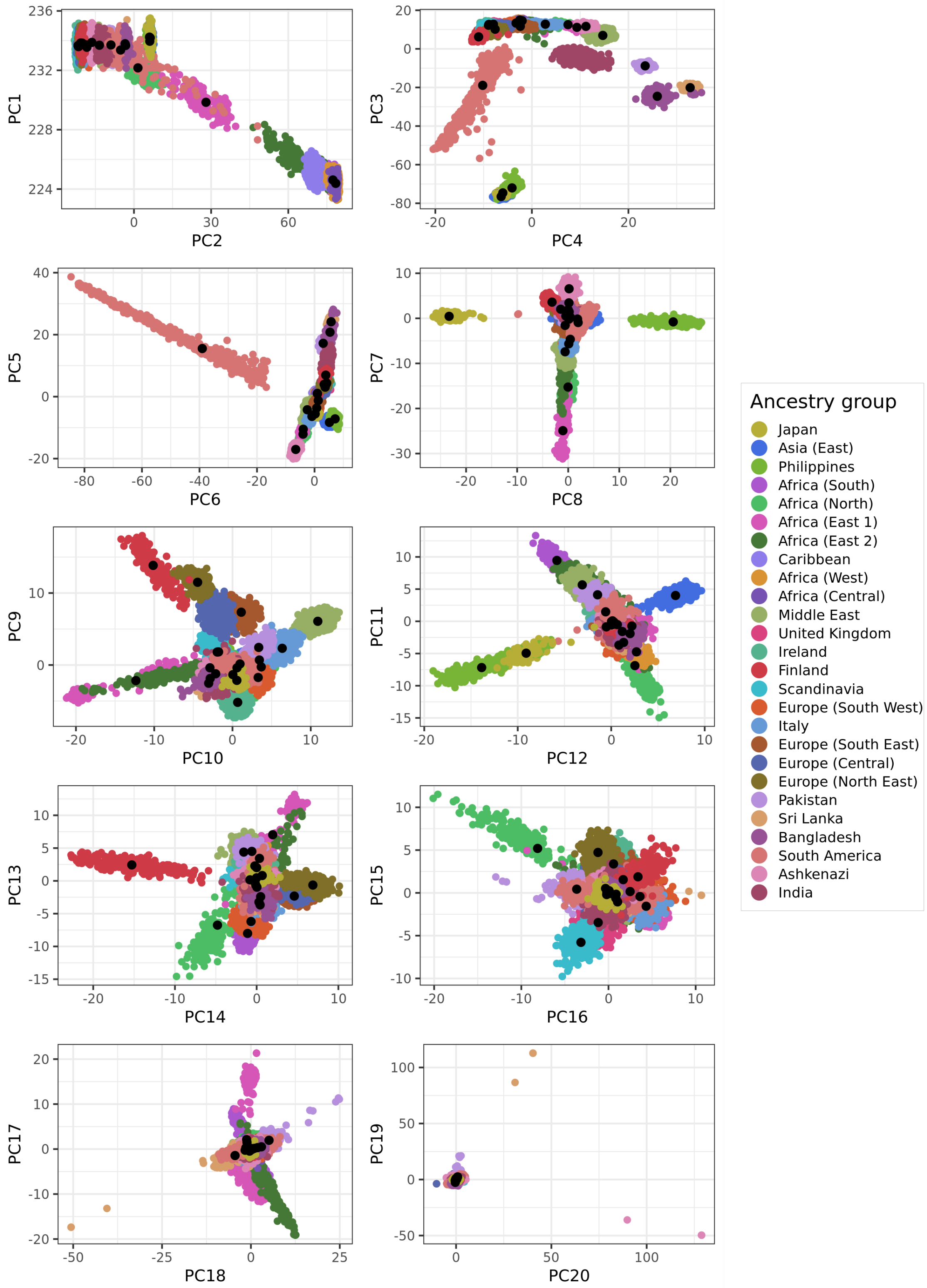


Figure S3: PCA scores for the PCA recomputed from the subset of UKBB and 1KG individuals used in the reference ancestry groups. Black points represent the projection of the allele frequencies from the 21 ancestry groups kept.

here), $K$ the number of reference populations (21 here), $p_l^{(k)}$ is the projection of frequencies from population $k$ onto PC $l$, and $p_l^{(0)}$ is the (corrected) projection of frequencies from the cohort of interest onto PC $l$. Note that we replace the quadratic term from this problem by the nearest positive-definite matrix to make it solvable, because of the lower number of PCs compared to populations. Note that, because they are very close ancestry groups, we merge a posteriori the ancestry coefficients $\alpha_k$ from "Ireland", "United Kingdom" and "Scandinavia" into a single "Europe (North West)" group, and similarly for "Europe (North East)" and "Europe (South East)" into a single "Europe (East)" group.

### S1.5 Supplementary results

| Ancestry group | N | BBJ | FinnGen | Perú | Qatar | Africa | GERA | PAGE | BrCa | PrCa | CAD | body fat | covid | eczema | epilepsy | urate |
|---|---|---|---|---|---|---|---|---|---|---|---|---|---|---|---|---|
| Africa (West) | 735 | 0.1 | | | | 34.8 | 2.1 | 26.4 | 0.2 | 0.2 | 2.4 | 0.5 | 4 | 0.4 | 0.7 | 3.2 |
| Africa (South) | 449 | | | | | 65 | 0.5 | 6.9 | 0.2 | 0.1 | 1.1 | 0.5 | 1.3 | 0.1 | 0.1 | 3.4 |
| Africa (East) | 276 | | | | 13 | 0.2 | | 0.3 | | | | 0.1 | 0.2 | | | 0.9 |
| Africa (North) | 268 | | | | 22.5 | | 0.1 | 1.8 | | | | | | | | |
| Middle East | 523 | | | | 60.8 | | 1 | | | 0.2 | 3 | 1.1 | 0.3 | | 0.8 | 1 |
| Ashkenazi | 1975 | | | | 1.5 | | 5.1 | 0.4 | 0.5 | 2.5 | 1.3 | 1.6 | 0.6 | 0.9 | 2.4 | 0.8 |
| Italy | 345 | | | | | | 5.1 | 2.4 | 2.7 | 3.3 | 3 | 3.8 | 0.8 | 3.4 | 5.6 | 2.5 |
| Europe (East) | 667 | | 7.6 | | | | 7.9 | 0.5 | 8.5 | 9.6 | 10.6 | 10.7 | 11.5 | 10.3 | 9.7 | 8.2 |
| Finland | 143 (+ 99) | | 83.6 | | | | 1.7 | 0.6 | 8.9 | 12.1 | 6.1 | 7.5 | 13 | 10 | 5.3 | 2.2 |
| Europe (North West) | 4416 | | 7.4 | | | | 57.8 | 9.5 | 67.7 | 63.7 | 51.7 | 57.2 | 60.8 | 69.4 | 66.4 | 43.8 |
| Europe (South West) | 603 | | | | | | 6.8 | 9 | 6 | 5.8 | 3.3 | 4.3 | 2.3 | 4.3 | 5.2 | 3.6 |
| South America | 473 (+ 84) | 0.2 | 0.6 | 98.2 | | | 4.6 | 24.9 | 1.1 | 0.7 | 1.4 | 0.5 | 1.9 | 0.5 | 0.3 | 0.1 |
| Sri Lanka | 372 | 0.1 | | | 2.2 | | | 0.8 | 0.7 | 0.1 | 2.1 | 1.6 | 0.3 | | | 0.2 |
| Pakistan | 400 | | | | | | | | 0.9 | 0.3 | 6.8 | 4.4 | 1.2 | | | 0.7 |
| Bangladesh | 223 (+ 86) | 0.2 | | | | | 0.3 | 1.3 | 1.4 | 0.9 | 2.9 | 2.4 | 0.9 | | | 0.7 |
| Asia (East) | 961 | 3.9 | | | | | 3.7 | 2.5 | 0.7 | 0.3 | 2.5 | 1.5 | 0.3 | 0.1 | 3.3 | 2.3 |
| Japan | 240 (+ 104) | 95.6 | 0.8 | 1.8 | | | 1.9 | 8.8 | 0.4 | 0.2 | 1.8 | 2.2 | 0.5 | 0.4 | | 26.2 |
| Philippines | 295 | | | | | | 1.4 | 4 | | | | | 0.2 | 0.1 | 0.1 | 0.2 |

Table S1: Reference populations with their size (N), and corresponding ancestry proportions (in %) inferred from the Summix method, for several GWAS summary statistics. Note that an equivalent of the Summix method was used, relying on a different quadratic programming solver, since Summix did not converge properly when using 21 reference populations. Note that, because they are very close ancestry groups, we merge a posteriori the ancestry coefficients $\alpha_k$ from "Ireland", "United Kingdom" and "Scandinavia" into a single "Europe (North West)" group, and similarly for "Europe (North East)" and "Europe (South East)" into a single "Europe (East)" group. Citations for the allele frequencies used are presented in Table 1 in the main text.

| Ancestry group | N | BBJ | FinnGen | Perú | Qatar | Africa | GERA | PAGE | BrCa | PrCa | CAD | body fat | covid | eczema | epilepsy | urate |
|---|---|---|---|---|---|---|---|---|---|---|---|---|---|---|---|---|
| Africa (West) | 735 | | | | | 31.3 | 1.8 | 27.6 | | | 3.2 | 0.9 | 3.9 | | 0.4 | 1.3 |
| Africa (South) | 449 | | | | | 68.7 | 1 | 5.9 | 0.4 | 0.4 | | 0.1 | 1.3 | 0.7 | | 4.4 |
| Africa (East) | 276 | | | | 12.1 | | | 0.1 | | | 0.2 | | 0.4 | | | 1.7 |
| Africa (North) | 268 | | | | 23.8 | | | | | | | | 0.3 | | 2.1 | |
| Middle East | 523 | | | | 63.7 | | | | | | 6.1 | 0.4 | | | 0.9 | 1.2 |
| Ashkenazi | 1975 | | | | | | 4 | 0.1 | | 2 | 0.6 | 0.7 | 0.7 | 0.5 | 2 | |
| Italy | 345 | | | | | | 3.8 | | 6.5 | 0.7 | | 8.5 | | | | 3.6 |
| Europe (East) | 667 | | | | | | 13.5 | 2.2 | 1.3 | 12.2 | 13.8 | 9 | 12.3 | 12.6 | 12.7 | 8.9 |
| Finland | 143 (+ 99) | | 99.9 | | | | 2.4 | | 9.2 | 13.1 | 6.9 | 8.8 | 13.6 | 14.9 | 3 | 3.7 |
| Europe (North West) | 4416 | | | | | | 57 | 4.1 | 67.5 | 63.9 | 54.5 | 60.3 | 62.2 | 67.9 | 65.5 | 44 |
| Europe (South West) | 603 | | | | | | 4.4 | 16.8 | 8.5 | 4.5 | | | | 2.5 | 9.9 | |
| South America | 473 (+ 84) | | | 100 | | | 4.5 | 25.6 | 1.4 | 0.7 | 1.5 | 0.5 | 1.6 | 0.3 | | |
| Sri Lanka | 372 | | | | 0.5 | | 0.3 | 2.4 | 3.6 | | 8.4 | 7 | 0.1 | | | 1.5 |
| Pakistan | 400 | | | | | | | | | | | | 2.5 | | | |
| Bangladesh | 223 (+ 86) | | | | | | | | | 1.9 | | | | | | |
| Asia (East) | 961 | | | | | | 3.5 | 3.1 | 1.1 | 0.6 | 2.2 | 1.1 | | 0.1 | 1.8 | |
| Japan | 240 (+ 104) | 100 | | | | | 2.1 | 8.3 | 0.4 | | 2.7 | 2.8 | 0.7 | 0.3 | 1.1 | 29.7 |
| Philippines | 295 | | 0.1 | | | | 1.6 | 3.7 | | | | | 0.5 | 0.3 | 0.5 | |

Table S2: Reference populations with their size (N), and corresponding ancestry proportions (in %) inferred from the proposed `snp_ancestry_summary` method while subsetting to 100,000 variants at random, for several GWAS summary statistics. Note that, because they are very close ancestry groups, we merge a posteriori the ancestry coefficients $\alpha_k$ from "Ireland", "United Kingdom" and "Scandinavia" into a single "Europe (North West)" group, and similarly for "Europe (North East)" and "Europe (South East)" into a single "Europe (East)" group. Citations for the allele frequencies used are presented in Table 1 in the main text.

double click on the icon to open the embedded Excel table

Table S3: Ancestry proportions (in %) for all 345 individuals of the Simons genome diversity project (Mallick *et al.* 2016) inferred either from the proposed `snp_ancestry_summary` method (using the genotypes of an individual divided by 2 in place of allele frequencies) or using the projection analysis of ADMIXTURE (`-P`, Shringarpure *et al.* (2016)). This table is also available in CSV format at `https://github.com/privefl/freq-ancestry/blob/main/ancestry_sgdp.csv`.
